# Maternal rejecting of newborns is epigenetic, intergenerationally transmitted and associated with altered miRNAs expression in owl monkeys

**DOI:** 10.1101/2025.07.01.662553

**Authors:** Jayde Farinha, Mirtha Clausi-Marroquin, Fabián Orlando Chamba Pardo, Nofre Sánchez-Perea, Charles A. Mein, Ping Yip, Ursula M. Paredes

**Affiliations:** Department of Anthropology, University College London, London, UK WC1H 0BW; Faculty of Veterinary Medicine, Universidad Nacional Mayor de San Marcos (UNMSM), Lima, Peru; Center for Technology Transfer, School of Veterinary Medicine, Universidad San Francisco de Quito, Ecuador; Center for Primate Conservation and Reproduction, Veterinary Institute of Tropical and High Altitude Research (IVITA-Iquitos), Faculty of Veterinary Medicine, Universidad Nacional Mayor de San Marcos (UNMSM), Lima, Peru; The Genome Centre, Blizard Institute, Queen Mary University of London, London, UK E1 2AT; Centre for Neuroscience, Surgery and Trauma, Blizard Institute, Queen Mary University of London, London, UK E1 2AT

**Keywords:** early-life adversity, intergenerational cycle, microRNAs, Aotus, owl monkey, epigenetic inheritance, captivity

## Abstract

Spontaneous maternal rejection of offspring is a considered a behavioural pathology affecting human and non-human primates, often with lethal consequences for newborns, yet it is widespread and pervasive across taxa, induced by stress or stress vulnerability. In some species, events of rejection can trigger chains of rejection which replicate themselves through generations. In this study we hypothesized that rejecting behaviours in owl monkeys are triggered and epigenetic in nature, transmitted to descendants, and their manifestation is associated with altered miRNA expression. Analysis of multi-generational records revealed that individuals who experienced rejection in infancy were 1.9 times more likely to reject their own offspring compared to controls, confirming its triggered nature. Transmission appears induced by experience rather than solely explained by genetic relatedness, as non-rejected full siblings showed no increased rejection tendency. Transcriptomic analysis identified distinct microRNA signatures associated with epigenetically induced rejecting behaviours. In rejected adults that become rejectors of their own offspring, mml-miR-1296, involved in lipid metabolism was significantly downregulated. In the naïve daughters of rejected-rejectors, upregulated mml-miR-125b-5p, previously identified as biomarker in humans exposed to early life trauma that also suffered behavioural pathology, was identified. In summary, these findings confirm the epigenetic nature of spontaneous rejecting, with robust intergenerational transmission, and accompanied by alteration of epigenetic marks across generations in owl monkeys.

## INTRODUCTION

Amongst captive non-human primates, it is not uncommon for newborns to experience maternal rejection or abandonment (Ryan et al., 2002, Prescott et al., 2012, Hopkins & Pearson 2000; Barnes & Cronin 2012, Sanchez et al., 2010). Without human intervention, rejected infants typically die (Johnson et al., 1991; Debyser 1995, Sanchez et al., 2006, Schino 2005, Fairbanks & McGuire 1995; Zipple et al., 2021) which brings a considerable number of rejected non-human primates to be hand or nursery reared (reviewed by Sackett et al., 2010). Individuals who experienced rejection in their early life and are nursery reared go on to develop behavioural disorders and often reject their own offspring (Porton & Niebruegge 2006, Suomi, 2005, Sproul-Bassett et al., 2020, Maestripieri, 2007). It has been documented that maternal rejection runs in non-human primate families in semi-captivity (Berman, 1990), suggesting a strong transmission of rejecting parental styles. Several mechanisms have been proposed to explain it. In nursery reared animals, transmission is often attributed to lack of adequate learning as infants grow without parents. In cross-fostering experiments of infant rhesus macaques with abusive mothers, it has been demonstrated that exposure to rejecting mothers is associated with increase rates of offspring neglect and abandonment in adopted daughters (Maestripieri, 2005). However, experiments also in rhesus macaques showed effect of this neglectful experience extends beyond behaviour. For example, monoamines and stress hormones concentration contribute to manifestation of rejecting behaviours, which are under genetic control (Maestripieri, 2008; Rogers, 2018). In recently studies in non-human primates, there were alterations in blood immune cells in primates that were parentally separated during early life, and surprisingly, effects were transmissible, and detectable across multiple generations (Kinnally & Capitanio, 2015, Kinnally et al., 2018). These observations suggest neglectful parental experience itself and genetic predisposition can be heritable.

Inheritance of acquired epigenetic marks has been proposed to explain transmission of experience through generations. In humans, experiencing neglect or rejection in mothers’ childhood is also associated with alterations in DNA methylation in genes of stress and neurodevelopment pathways, alterations in genes detail detectable in their children (León et al., 2022, Yehuda et al., 2016). In animals small non-coding RNAs also participate in transmission of experience through generations (Gapp et al., 2014, Jawaid 2018; Cecere, 2021, Rechavi & Lev, 2017). One hypothesis proposes that in mammals, small non-coding RNAs produced in central nervous system are circulatory by entering biofluids then reaching germline (Devanapally et al., 2015; Miska & Fergusson-Smith 2016; Jawaid et al., 2018), therefore explaining how stress experience can pass down to next generations. Within these, microRNAs are small epigenetic modifiers that regulate gene expression (Kinser and Pincus 2020). These molecules can regulate the transcriptome through the RNA-induced silencing complex (RISC) and binding to the 3’ untranslated region (UTR) of their target transcripts, resulting in reduction of mRNA expression (Smith-Vikos & Slack, 2012). In parallel, genetic epidemiological studies showed that in adults exposed to early life stress and experienced altered behavioural pathologies in adulthood, circulatory miRNA expression is altered, associated to differences in DNA methylation (Cattane et al., 2019; Cattaneo et al., 2020). In sperm of adults and serum of children exposed to childhood trauma and neglect, altered miRNAs in plasma and germline are observed (Gomółka et al., 2023). It is possible that experienced triggered miRNAs participate in the transmission of spontaneous rejecting behaviours in captive non-human primates. Studying this fascinating phenomenon could help identify factors that contribute to rejection transmission, a significant problem for their welfare in captivity.

Our previous work has characterised maternal rejection in a captive population of owl monkeys *Aotus nancymaae* where rejections occur spontaneously (Sanchez et al., 2006). Owl monkeys belong to a genus composed of monogamous species that display biparental care (Fernandez-Duque, 2012).We have previously shown that rejected owl monkey infants, despite being rescued by veterinarians, show persistent hormonal alterations (Osman et al., 2021), accelerated epigenetic ageing (Girling et al., 2024), and if they survive to adulthood, a reduced life expectancy and fitness, and behaviourally elevated stress reactivity and rejectfulness towards their young, and effects detectable in the following generation (Farinha & Clausi et al., 2024). Additionally, daughters of rejected individuals show upregulation of specific miRNAs associated with stress-related disorders in humans (Cattane et al., 2020), demonstrating miRNAs are associated in the transmission of intergenerational effects of rejection.

One effect not studied in this population is the transmission of rejecting behaviours themselves. The present report tackles this question. First, using multigenerational, individual-based population records of we test: i) whether being rejected trigger being rejectful towards young in adulthood, ii) if rejecting behaviours increase in the offspring of rejected, iii) whether rejecting transmission is explained by genetic or epigenetic roots, and iv) if transmission is associated with altered miRNA signatures across generations.

## MATERIALS AND METHODS

### Ethics

In this study, we reanalyzed data from colonies maintained at the Center for Reproduction and Conservation of Non-Human Primates (CRCP) of IVITA at the National University of San Marcos (UNMSM), located in Iquitos, Peru.

### Housing

The animals were housed under standardized conditions in individual enclosures (2 m^3^) with access to natural biological periods and fed a diet of seasonal fruits, balanced dry food, and water ad libitum.

### Demographic analyses

To examine the transgenerational transmission of rejecting behaviour, we conducted a series of logistic regression analyses on a dataset of 736 individuals. These analyses aimed to determine whether experiencing parental rejection in one generation influenced the likelihood of exhibiting rejection behaviours in the next. Three primary models were tested:

### 1. Direct intergenerational transmission

We examined whether individuals who were themselves rejected as infants were more likely to reject their own offspring, indicating a direct behavioural transmission across generations.

### 2. Sibling analysis

We assessed whether non-rejected siblings of rejected individuals exhibited an elevated likelihood of rejecting their own offspring, suggesting indirect familial influences or shared environmental/epigenetic effects.

### 3. Offspring of rejected

We investigated whether the offspring of parents who were rejected as infants were more likely to become rejectors themselves, reflecting potential vertical transmission of behaviour through both biological and social mechanisms.

All models included relevant covariates where appropriate (e.g., sex or cohort effects), and rejection behaviour was modelled as a binary outcome (rejected vs. not rejected). Confidence intervals and predicted probabilities were extracted to interpret the magnitude and significance of effects.

**RNA sequencing analyses**, we reanalyzed data generated in a previous experiment characterizing miRNA profiles of rejected owl monkeys (Farinha and Clausi 2024). The original study collected blood samples under sedation (ketamine 8 mg/kg) via femoral venipuncture, with 0.5 ml per primate placed on filter paper. Total RNA was extracted from dried blood spots (3 × 6mm punches per owl monkey, n=48) using the mirVana miRNA isolation kit (Thermo Fisher Scientific) as previously described (Ponnusamy et al., 2021), concentrated using the RNeasy MinElute kit (Qiagen), and libraries were prepared using the QIAseq miRNA library kit. Sequencing was performed on a NextSeq 2000 platform (Illumina Inc.).

**RNA sequencing processing and analysis** Protocol was detailed in previously described (Farinha & Clausi et al., 2024). In brief, sequencing reads were trimmed using Trimgalore v0.6.6 and aligned to the owl monkey genome (*Aotus nancymaae*, Anan_2.0) using STAR v2.7.3a. Gene counts were summarized using HTseq and analyzed in Partek Flow. RNA results were mapped against *Aotus nancymaae* and *Macaca mulatta* genomes for miRNA identification. Differential expression analysis was performed using LIMMA TREND with Benjamini-Hochberg FDR correction (FDR<0.1) and a ±1.5 cutoff value.

To examine intergenerational transmission of rejection, we compared individuals who experienced parental rejection and became rejectors themselves (rejected-rejectors, n=4) with those who were rejected but did not become rejectors (rejected-only, n=4) (Figure 1). Additionally, we analyzed miRNA expression patterns in the well-reared offspring of these groups (offspring of rejected-non rejectors: n=7; offspring of rejected-rejectors: n=5).

**Figure 1.**
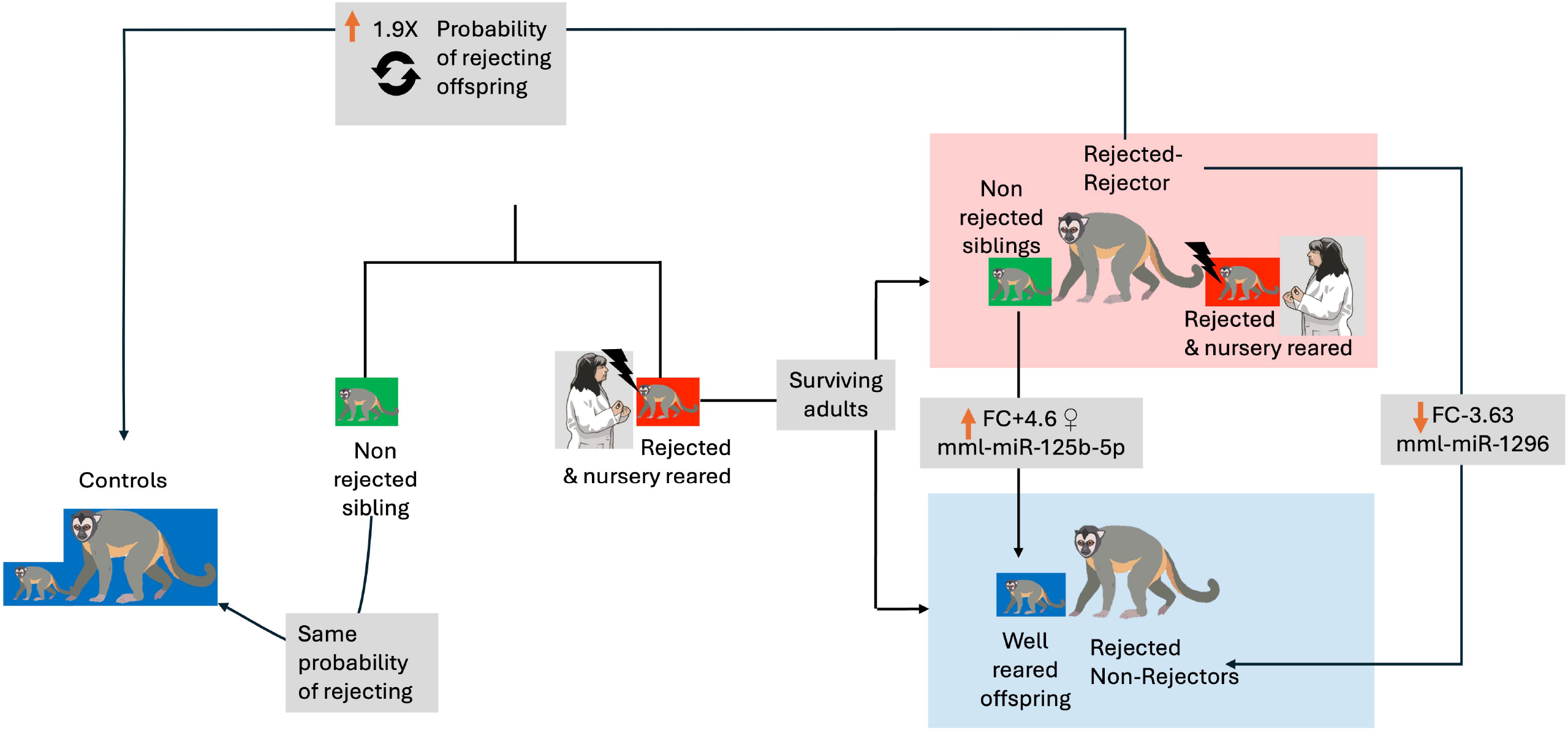
Graphical Abstract depicting the cyclical transmission of rejecting behaviors in owl monkeys. Control animals (dark blue boxes) represent normally reared individuals, while rejected infants are rescued and nursery reared (red boxes). Surviving rejected adults show a 1.9-fold increased probability of rejecting their own offspring compared to controls. This transmission is experience-dependent rather than genetic, as non-rejected siblings (green boxes) show the same rejecting probability as controls. The molecular basis of rejecting transmission involves distinct microRNA signatures. Rejected individuals who become rejectors in adulthood (pink boxes) show downregulation of mml-miR-1296 (fold change = -3.63), a microRNA involved in metabolism. Their well-reared offspring (light blue boxes) display striking upregulation of mml-miR-125b-5p with a 4.6-fold increase, particularly in daughters (♀). This microRNA is a known biomarker of early life trauma in humans, suggesting a conserved epigenetic response that transmits across generations and predisposes offspring to similar behavioural patterns.

## RESULTS

### Rejection behaviors are transmissible and occur more frequently in individuals with a history of early-life rejection

We conducted a logistic regression to determine whether individuals with a history of parental rejection were more likely to reject their own offspring. The coefficient for individuals with parental rejection history versus those without (β1 = 0.64) was statistically significant (p < 0.01), indicating that being rejected is associated with increased log odds (1.9 times higher) of becoming a rejector (Figure 2a). The results suggest that individuals with a history of parental rejection have higher odds of exhibiting rejection behaviours towards their own offspring (p = 0.034, also Figure 2a).

**Figure 2.**
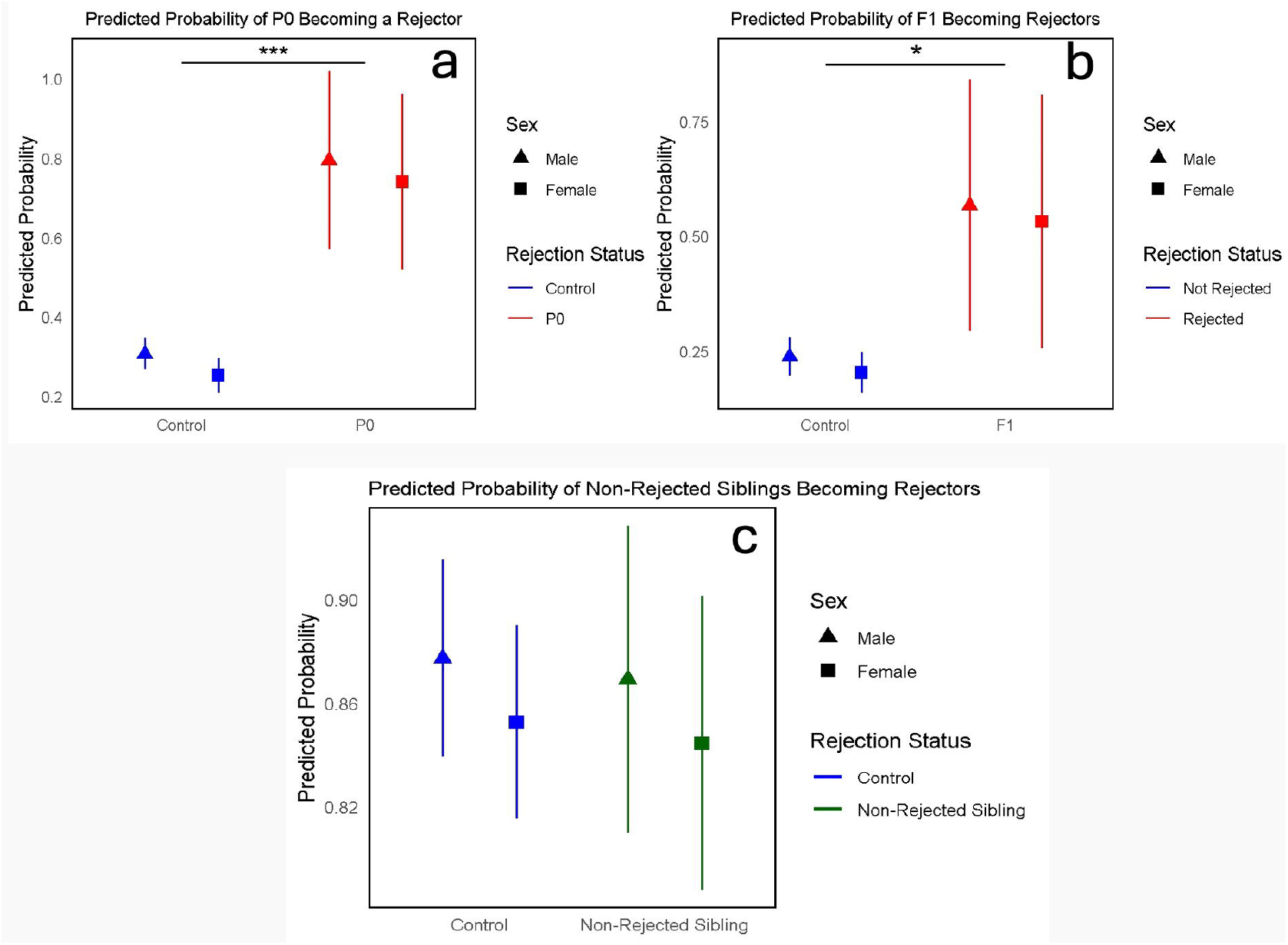
Diagram depicting the rejecting transmission cycle in owl monkeys. 2. Probability of rejecting newborns in rejected owl monkeys, their siblings and offspring. a. Probability of rejected on becoming rejectors versus controls, b. Probability of offspring of rejected (Fl) of becoming rejectors is increased versus controls, c. Probability of becoming rejectors in well-reared siblings of rejected is the same as for controls. Asterisks indicate statistically significant differences between groups, where ^*^= p<0.05, where and ^***^= p<0.005.

The log odds of rejecting a newborn offspring increased significantly by two-fold when one has a parent who has been rejected (Figure 2b). In contrast, the likelihood of becoming a rejector in well reared siblings of rejected owl monkeys was similar to controls (β1 = 0.02551, p = 0.1002, Figure 2c). This indicates that “inherited” rejection behaviours are potentially conditional upon the experience or physical and physiological conditions associated with rejection in offspring, rather than genetic ancestry alone.

### Rejecting behaviours in rejected animals are associated with dysregulation of microRNAs in blood across two generations

As rejecting behaviours were more frequent in offspring of rejected individuals, we performed differential expression analysis of whole blood miRNAs in animals that reject their offspring, and between rejected individuals who either reject their own offspring or not, and their naive descendants. Overall, rejected in infancy who become rejectors in adulthood showed distinct differences from rejected who did not reject. We identified 25 miRNAs significantly altered, with 3 downregulated and 22 upregulated (p<0.05, Figure 3.a,) Specifically, miR-1296 was downregulated in rejected-rejectors (FDR-p = 0.05, fold change = -3.63, indicated by asterisk).

**Figure 3.**
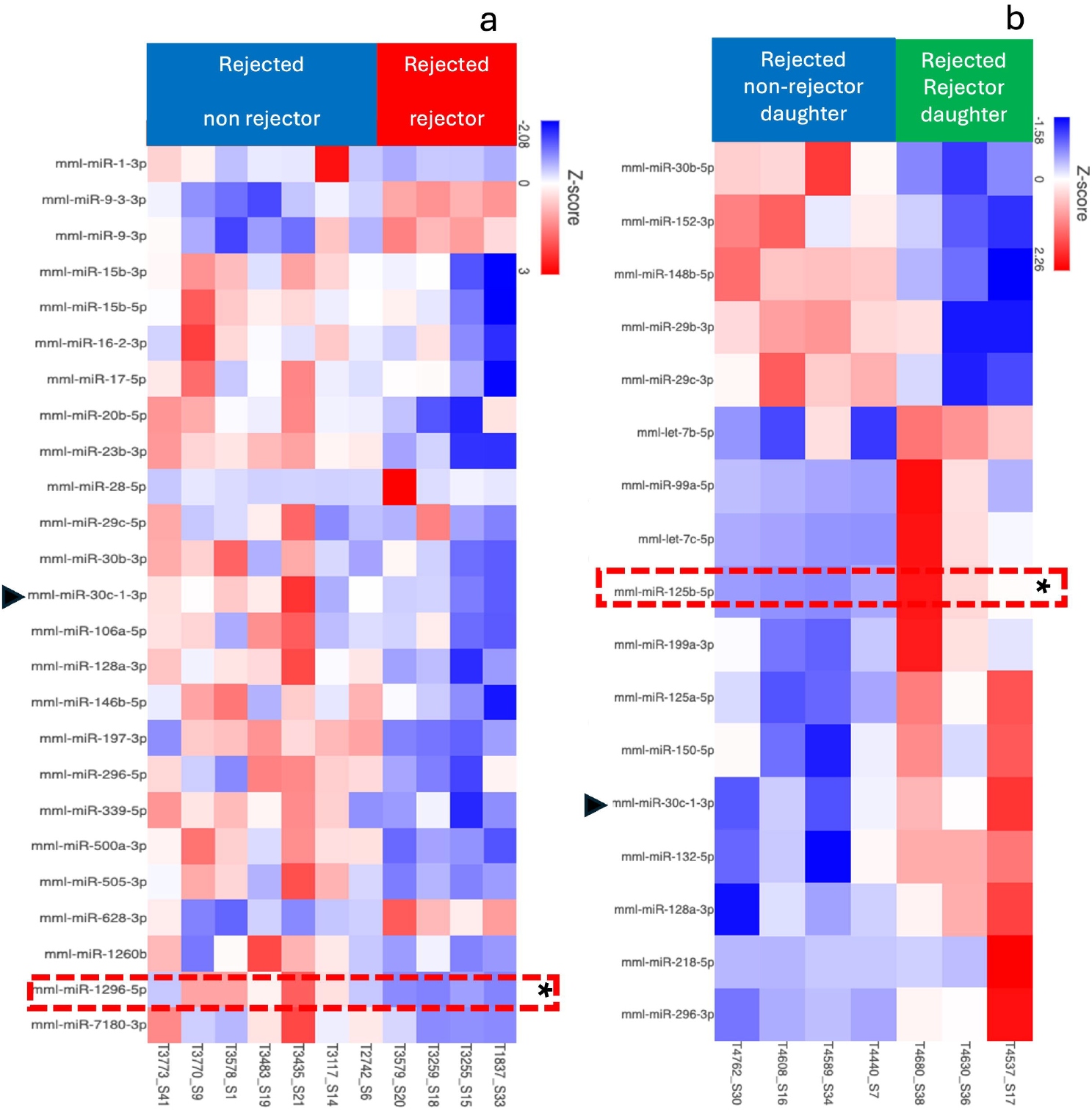
Rejecting behaviours in rejected animals and descendants were associated with dysregulation of microRNAs in blood. (a)Hcatmap with standardised Z-scores showed 25 miRNAs that were dysrcgulatcd between rejected and rejectors, (b) Heatmap with standardised Z-scores showed 17 miRNAs were dysregulated between their naive daughters (p<0.05). Red blocs indicate upregulated and blue blocks indicate downregulation. A black asterisk indicates miRNAs which p value passed FDR threshold. Arrow heads indicate miRNAs that were dysregulated in two generations.

A molecular signature of rejecting behaviours were detected in naive offspring (Figure 3.b), particularly in females. Overall, offspring of rejected-rejectors showed increased expression of 12 upregulated, and 5 downregulated (p<0.05), but only mml-miR-125b-5p (indicated by asterisk) passed multiple comparison threshold (2.24-fold increase, FDR-p = 0.044). This effect was particularly pronounced in daughters of rejected-rejectors displaying a striking 4.6-fold increase when compared to daughters of rejected non rejectors (daughters of rejected rejectors (n=3), of rejected non rejectors n=4, FDR-p = 0.002). Male offspring showed some miRNA changes with a p value<0.05, but none of these changes reached statistical significance after controlling for multiple comparisons. One miRNA was seen altered in both generations mml-miR-30c-1-3p although with opposite expression (black arrow heads, Figure 3, left and right panels), downregulated in rejected-rejectors mothers (FC=-5 rejected-rejectors/rejected-non rejectors mothers) and upregulated in their daughters (FC=2 rejected-rejector/rejected non-rejector daughters).

Two other microRNAs were seen altered in parents and offspring, however, they displayed arm switch expression: mml-miR-29c-5p and mml-30b-5p were both found downregulated in rejected-rejectors, and the opposite arm in the precursor miRNA was downregulated in their daughters.

## DISCUSSION

Our study provides compelling evidence for the intergenerational transmission of rejection behaviours in owl monkeys. This experience-dependent transmission is heritable to offspring. At the molecular level, specific microRNA expression signatures are associated with rejecting behaviours, with particularly pronounced effects in daughters, suggest possible involvement of miRNAs in rejectors.

Owl monkeys of both sexes that were rejected have a reduced chances of survival to reproductive age (Farinha & Clausi et al., 2024). If they survive to reproductive age and bare offspring, they have an increased likelihood of rejecting their own young newborns, and even their offspring also show increased odds of rejecting behaviours. This present study proposes the second intergenerational effect associated with rejection experience detected in this owl monkey population and joins other studies demonstrating transmission of neglect in mammals (in rodents: Curley et al., 2008; non-human primates: Sproul-Bassett et al., 2020; Maestripieri 2005; Berman 1990 and humans: Brockington, 2011; Gilbert & Lacey, 2021).

Rejecting behaviour expression and transmission in this colony appear epigenetic, as their incidence is linked to experiencing rejection rather than be attributable to genetic ancestry alone. Our study agrees with previous landmark studies using cross-fostering on non-human primates, showing that infants reared by abusive rhesus macaque mothers (whether biological or adoptive) also abused their infants, showing the intergenerational transmission of infant abuse was not explain by genetic inheritance but rather induced by experience (Maestripieri, 2005). More recent epidemiologically studies in humans demonstrate a link between behavioural and epigenetic mechanisms of neglect transmission. These studies showed that differences in DNA methylation profiles, shared between neglectful mothers and their offspring and linked to adverse experiences of mothers in early life (León et al., 2022). In a follow up study, these authors developed an epigenetic maternal neglect score, where greater epigenetic burden was identified in mothers who having experienced neglect were also neglectful towards their own offspring (León et al., 2025).

However, multiple exposures other than epigenetics have been suggested to explain transmission of rejecting behaviours. In non-human primates, poor maternal condition has been proposed as a significant contributor to caregiving quality and offspring wellbeing. For example, in vervet monkeys, mothers in marginal conditions exhibit higher rates of maternal rejection compared to those in average condition (Fairbanks & McGuire, 1995). A high-fibre diet intervention to improve food quality in captive vervet monkey led to intended weight loss and also unintended increases in maternal rejection, particularly in lower-weight mothers (Fairbanks et al., 2010), In a cross taxa review of maternal death and effects on offspring survival showed that mother’s poor health render mothers less capable to provide adequate care for their young (Zipple et al., 2021).

Accordingly, we have previously shown that poor condition is associated with experience of rejection in owl monkeys. We reported increased visits to clinics and elevated stress in rejected infants (Osman et al., 2021), lifetime accelerated epigenetic ageing (Girling et al., 2024), poor reproductive outcomes and early death, and infant behavioural rejection and increased stress reactivity, effects also detectable in their offspring (Farinha & Clausi 2024). The rates of rejecting within the rejected, and their descendants are also significantly higher. As they co-occur, it is possible that poor condition might indeed contribute to increase rates of rejecting tendencies in owl monkey mothers. Some mothers who were rejected in early life that go on to become rejectors reject multiple times (Supplementary material 1), and some of their offspring live long lifespans (Supplementary material 2), demonstrating good condition. Further, health was monitored daily across this colony, and no apparent differences reported in health records were identifiable in rejecting and non-rejecting mothers (N. Sanchez, personal communication 30^th^ of June 2025). Whilst we previously documented transmission of poor condition, stress reactivity and of poor parental care, these does not fully explain why some rejected owl monkey mothers become rejectors.

Learning neglectful maternal styles is often cited as a possible explanation for intergenerational transmission of neglect. Undeniably, rejected and nursery reared owl monkeys have not been exposed to adequate parenting in early life, likely leaving a biobehavioural sequela in the rejected (Pryce et al., 2004, Sanchez et al., 2006; Sackett et al., 2010). However, in the present study, incidences of rejection mostly occur in first births and without precedent (Supplementary material 1), successfully rearing some infants to adulthood, demonstrating adequate rearing skills. Therefore, transmission of rejecting from one generation to the next in these owl monkeys is not solely explained by lack of learning opportunities or skill in those rejected and nursery reared.

Antecedents of epigenetic transmission of newborn rejecting do not exist in non-human primates. However, there is evidence of induced stress experience being transmitted through non-genetic mechanisms in macaques. Maternal separation studies in rhesus macaques showed increased nervousness (after 3 generations) and reduced white blood cell counts (after 2 generations) in descendants of male juveniles that were nursery reared in infancy (Kinnally et al., 2018). Stress reactivity is also a risk factors for infant neglect in young primate mothers (Suomi 1996) and multiple studies demonstrate that parental separation triggers variation in maternal behaviours and epigenetic marks, paralleling our observations in owl monkeys (stress reactivity and serotonergic function: Kinnally et al., 2010; bonding: oxytocin receptor genes Baker et al., 2017). Then it is plausible that transmission of rejecting in owl monkeys is mediated by epigenetic modifications in genes involved in stress vulnerability and bonding.

We targeted microRNAs in blood, to characterise distinguishing epigenetic features of rejected rejectors parents and their descent. MicroRNAs are capable to pass the Weisman barrier, a separation between germline cells and somatic cells (Eaton et al., 2015). If miRNAs were induced by maternal rejection, these could be expressed in somatic cells, become circulatory and then reach germline, then passing down through generations (Jawaid et al., 2018, Miska & Fergusson-Smith, 2016, Rechavi & Lev, 2017).

Our study identified one microRNA (mml-miR-1296) being significantly down-regulated (mml-miR-1296) in rejected-rejectors. The function of this miRNA in owl monkeys is unknown, in humans its altered expression has been associated with metabolic dysregulation. Down-regulation of miR-1296 has also been identified as a biomarker for non-alcoholic fatty liver disease (NAFLD) (Yu et al., 2019). Also, this miRNA participates in the metabolic adaptation enabling lactation transition in dairy cows (Veshkini et al., 2022) and is altered in plasma exosomes of premature human infants fed formula compared to those fed breast milk (Kim et al., 2024). Anecdotally, newborn rejected owl monkeys are fed formula, and some recorded events of rejection occur when mothers can not or make little milk (N. Sanchez, 30^th^ of June, 2025).

Further, our analysis showed alterations of miR-125b-5p, a conserved feature of early life adversity response, in female offspring of rejected that became rejectors themselves. In humans, has-miR-125b is preferentially present in the brain, and female reproductive system (GTEx Gene VB track, Assembly Hg38, **http://genome.ucsc.edu**). Its roles are implicated in cardiac development and protection against ischemic injury (Chen et al., 2021). In adults, miR-125b-1-3p (same precursor as miR-125b-5p) was found over-expressed in blood of adults with a history of childhood trauma, where overexpression confers increased risk of developing schizophrenia (Cattane et al., 2019). In the same study it was shown that rats exposed to prenatal stress induced the overexpression of this miRNA in the hippocampus. Overexpressed miR-125b contributes to regulated cortisol metabolism (Robertson et al., 2017), produces anxiety like behaviour in mice (Chen et al., 2024) and is expressed peripherally in saliva of children exposed to early life trauma and parental loss (Ozaki et al., 2022). Conversely, miR-125b can be induced and reported as protective against oxidative stress, through regulation of the transcription factor p53 (Le et al., 2009). Also, its overexpression is considered protective against UVB-induced accelerated ageing (Cui et al., 2025), and against acidosis-induced neuronal death in a rat cerebral stroke model (Dong et al., 2024).

Finally, we explored other altered miRNAs dysregulated (p<0.05) which did not pass threshold FDR to avoid neglecting relevant biological phenomenon. This approach highlighted mml-mR-30c-1-3p, which altered in both generations but expressed in opposite direction in parents and offspring (Figure 3). This miRNA is one shown to be intergenerationally inherited and has been associated in transmission of altered hormonal synthesis in response to stress in cadmium-exposed pregnant rats (Luo et al., 2023) and implicated in neurodegenerative and cerebrovascular disorders (Kumar and Li 2022).

As no miRNA was altered in both generations which passed FDR threshold, our study suggests no detection of intergenerational transmission of circulatory miRNAs in blood. However, as both miRNAs have developmentally, and tissue specific expression patterns and our sample size was small, whether miRNAs induced by rejection are intergenerationally transmitted require further research. Nonetheless, the results do suggest that alterations in miRNA candidates in offspring linked to stress and early life trauma and for lipid and energy metabolism in parents, in agreement with previous findings Similarly, we found sex specific effect, also in line with previously reported (e.g. Kinnally et al 2018, Boscardin et al., 2022).

Newborn rejecting transmission is an evolutionary paradox given the profound negative consequences on fitness in owl monkeys (Farinha & Clausi, 2024). Previous observations of spontaneous rejecting in the wild and captivity proposed that maternal rejection and neglect might represent and extreme form of adaptive recalibration of energetic investment under stressful conditions (e.g. Melo et al., 2003, Culot et al., 2011 Maestripieri & Carroll, 2000). Trivers (1974) established that parents incur into high metabolic costs which are balanced against survival of present and future offspring reproduction. If indeed rejecting might is an extreme trade-off, its transmission, mediated by induced epigenetic alterations may be adaptive for primate mothers facing poor resource or stressful environments, and the very few offspring which survive. Incidental finding that extended lifespans are associated with rejecting phenotypes warrants future studies to test this intriguing hypothesis in the owl monkeys study system.

In conclusion, in this owl monkey colony, rejecting and intergenerational transmission in owl monkeys does occur, this is triggered by experience, multifactorial and epigenetic.

Further, overexpression of mml-miR-125b and mml-miR-1296 and possibly mml-miR-30c-1-3p are associated with altered maternal phenotypes and parental adverse experience. As these are found exclusively in the rejected-rejector and their descendants, our results suggest that miRNAs are not just associated with negative intergenerational effects of rejection (Farinha & Clausi 2024) but also with intergenerational cycles of rejecting in owl monkeys. If indeed altered miRNA were functional and physiologically relevant in owl monkeys, differences in stress vulnerability and metabolism might distinguish future rejectors, and their offspring, and provide a therapeutic target.

## REFERENCES

1. Baker M, Lindell SG, Driscoll CA, Zhou Z, Yuan Q, Schwandt ML, Miller-Crews I, Simpson EA, Paukner A, Ferrari PF, Sindhu RK, Razaqyar M, Sommer WH, Lopez JF, Thompson RC, Goldman D, Heilig M, Higley JD, Suomi SJ, Barr CS. Early rearing history influences oxytocin receptor epigenetic regulation in rhesus macaques. Proc Natl Acad Sci U S A. 2017 Oct 31;114(44):11769–11774.

2. Barnes HA, Cronin A. Hand‐rearing and reintroduction of Woolly monkey Lagothrix lagotricha at Monkey World–A pe Rescue Centre, UK. International Zoo Yearbook. 2012 Jan;46(1):164–74.

3. Berman CM. Intergenerational transmission of maternal rejection rates among free-ranging rhesus monkeys. Animal Behaviour. 1990 Feb 1;39(2):329–37.

4. Bohacek J, Mansuy IM. Molecular insights into transgenerational non-genetic inheritance of acquired behaviours. Nature Reviews Genetics. 2015 Nov;16(11):641–52.

5. Boscardin C, Manuella F, Mansuy IM. Paternal transmission of behavioural and metabolic traits induced by postnatal stress to the 5th generation in mice. Environ Epigenet. 2022 Nov 17;8(1):dvac024.

6. Brockington I. Maternal rejection of the young child: present status of the clinical syndrome. Psychopathology. 2011 Jul 7;44(5):329–36.

7. Cattane N, Mora C, Lopizzo N, Borsini A, Maj C, Pedrini L, Rossi R, Riva MA, Pariante CM, Cattaneo A. Identification of a miRNAs signature associated with exposure to stress early in life and enhanced vulnerability for schizophrenia: New insights for the key role of miR-125b-1-3p in neurodevelopmental processes. Schizophrenia research. 2019 Mar 1;205:63–75.

8. Cattaneo A, Suderman M, Cattane N, Mazzelli M, Begni V, Maj C, D’Aprile I, Pariante CM, Luoni A, Berry A, Wurst K. Long-term effects of stress early in life on microRNA-30a and its network: Preventive effects of lurasidone and potential implications for depression vulnerability. Neurobiology of Stress. 2020 Nov 1;13:100271.

9. Cecere G. Small RNAs in epigenetic inheritance: from mechanisms to trait transmission. FEBS Lett. 2021 Dec;595(24):2953–2977.

10. Chen CY, Lee DS, Choong OK, Chang SK, Hsu T, Nicholson MW, Liu LW, Lin PJ, Ruan SC, Lin SW, Hu CY, Hsieh PCH. Cardiac-specific microRNA-125b deficiency induces perinatal death and cardiac hypertrophy. Sci Rep. 2021 Jan 27;11(1):2377.

11. Chen H, Wu J, Zhu X, Ma Y, Li Z, Lu L, Aschner M, Su P, Luo W. Manganese-induced miR-125b-2-3p promotes anxiety-like behavior via TFR1-mediated ferroptosis. Environmental Pollution. 2024 Mar 1;344:123255.

12. Cui H, Fu LQ, Teng Y, He JJ, Shen YY, Bian Q, Wang TZ, Wang MX, Pang XW, Lin ZW, Zhu MG. Human Hair Follicle Mesenchymal Stem Cell-Derived Exosomes Attenuate UVB-Induced Photoaging via the miR-125b-5p/TGF-β1/Smad Axis. Biomaterials Research. 2025 Jan 13;29:0121.

13. Culot L, Lledo-Ferrer Y, Hoelscher O, Muñoz Lazo FJ, Huynen MC, Heymann EW. Reproductive failure, possible maternal infanticide, and cannibalism in wild moustached tamarins, Saguinus mystax. Primates. 2011 Apr;52:179–86.

14. Curley JP, Champagne FA, Bateson P, Keverne EB. Transgenerational effects of impaired maternal care on behaviour of offspring and grandoffspring. Animal Behaviour. 2008 Apr 1;75(4):1551–61.

15. Debyser IW. Platyrrhine juvenile mortality in captivity and in the wild. International journal of primatology. 1995 Dec;16:909–33.

16. Devanapally S, Ravikumar S, Jose AM. Double-stranded RNA made in C. elegans neurons can enter the germline and cause transgenerational gene silencing. Proceedings of the National Academy of Sciences. 2015 Feb 17;112(7):2133–8.

17. Dong K, Chen F, Wang L, Lin C, Ying M, Li B, Huang T, Wang S. iMSC exosome delivers hsa-mir-125b-5p and strengthens acidosis resilience through suppression of ASIC1 protein in cerebral ischemia-reperfusion. Journal of Biological Chemistry. 2024 Aug 1;300(8).

18. Eaton, S. A., Jayasooriah, N., Buckland, M. E., Martin, D. I., Cropley, J. E., & Suter, C. M. (2015). Roll Over Weismann: Extracellular Vesicles in the Transgenerational Transmission of Environmental Effects. Epigenomics, 7(7), 1165–1171.

19. Fairbanks LA, McGuire MT. Maternal condition and the quality of maternal care in vervet monkeys. Behaviour. 1995 Aug 1:733–54.

20. Fairbanks LA, Blau K, Jorgensen MJ. High‐fiber diet promotes weight loss and affects maternal behavior in vervet monkeys. American Journal of Primatology: Official Journal of the American Society of Primatologists. 2010 Mar;72(3):234–41.

21. Farinha J, Clausi M, Chamba Pardo FO, Sanchez-Perea N, Boot J, Mein CA, Yip P, Paredes UM. Parental rejection is associated with intergenerational inheritance of reduced survival, altered behaviours and miRNAs in Owl Monkeys. bioRxiv. 2024 Dec 20:2024–12.

22. Fernandez‐Duque E. Owl monkeys Aotus spp in the wild and in captivity. International Zoo Yearbook. 2012 Jan;46(1):80–94.

23. Franklin TB, Russig H, Weiss IC, Gräff J, Linder N, Michalon A, Vizi S, Mansuy IM. Epigenetic transmission of the impact of early stress across generations. Biological psychiatry. 2010 Sep 1;68(5):408–15.

24. Gapp K, Jawaid A, Sarkies P, Bohacek J, Pelczar P, Prados J, Farinelli L, Miska E, Mansuy IM. Implication of sperm RNAs in transgenerational inheritance of the effects of early trauma in mice. Nature neuroscience. 2014 May;17(5):667–9.

25. Gilbert R, Lacey R. Intergenerational transmission of child maltreatment. Lancet Publ Health. 2021;6:e435–6.

26. Girling MT, Sanchez NM, Paredes UM. Global DNA Hypomethylation as a Biomarker of Accelerated Epigenetic Ageing in Primates. bioRxiv. 2024 Mar 30:2024–03.

27. Gomółka M, Tomaszewska W, Canalda AJ, Kamińska-Kaczmarek B, Jawaid A. Distinct microRNA signatures of childhood trauma in human serum and sperm: Implications for potential intergenerational transmission. IBRO Neuroscience Reports. 2023 Oct 1;15:S924–5.

28. Hopkins WD, Pearson K. Chimpanzee (Pan troglodytes) handedness: variability across multiple measures of hand use. Journal of Comparative Psychology. 2000 Jun;114(2):126.

29. Jawaid A, Roszkowski M, Mansuy IM. Transgenerational Epigenetics of Traumatic Stress. Prog Mol Biol Transl Sci. 2018;158:273–298.

30. Johnson LD, Petto AJ, Sehgal PK. Factors in the rejection and survival of captive cotton top tamarins (Saguinus oedipus). American Journal of Primatology. 1991;25(2):91–102.

31. Kim EB, Song JH, L. LN, Kim H, Koh JW, Seo Y, Jeong HR, Kim HT, Ryu S. Characterization of exosomal microRNAs in preterm infants fed with breast milk and infant formula. Frontiers in Nutrition. 2024 Jan 18;11:1339919.

32. Kinnally EL, Capitanio JP, Leibel R, Deng L, LeDuc C, Haghighi F, Mann JJ. Epigenetic regulation of serotonin transporter expression and behavior in infant rhesus macaques. Genes, Brain and Behavior. 2010 Aug;9(6):575–82.

33. Kinnally EL, Capitanio JP. Paternal early experiences influence infant development through non-social mechanisms in Rhesus Macaques. Frontiers in Zoology. 2015 Dec;12:1–8.

34. Kinnally EL, Gonzalez MN, Capitanio JP. Paternal line effects of early experiences persist across three generations in rhesus macaques. Developmental Psychobiology. 2018 Dec;60(8):879–88.

35. Kinser HE, Pincus Z. MicroRNAs as modulators of longevity and the aging process. Human genetics. 2020 Mar;139(3):291–308.

36. Kumar M, Li G. Emerging Role of MicroRNA-30c in Neurological Disorders. Int J Mol Sci. 2022 Dec 20;24(1):37.

37. Le MT, Teh C, Shyh-Chang N, Xie H, Zhou B, Korzh V, Lodish HF, Lim B. MicroRNA-125b is a novel negative regulator of p53. Genes & development. 2009 Apr 1;23(7):862–76.

38. León I, Herrero Roldán S, Rodrigo MJ, López Rodríguez M, Fisher J, Mitchell C, Lage-Castellanos A. The shared mother-child epigenetic signature of neglect is related to maternal adverse events. Front Physiol. 2022 Aug 24;13:966740.

39. León, I., Góngora, D., Rodrigo, M.J. et al. Maternal epigenetic index links early neglect to later neglectful care and other psychopathological, cognitive, and bonding effects. Clin Epigenet 17, 46 (2025).

40. Luo L, Li J, Sun Y, Lv Y, Liu J, Li Y, Zhang C, Zhang W. Maternal genetic intergenerational and transgenerational effects on hormone synthesis in ovarian granulosa cells of offspring exposed to cadmium during pregnancy. Ecotoxicology and Environmental Safety. 2023 Sep 15;263:115278.

41. Maestripieri D, Carroll KA. Causes and consequences of infant abuse and neglect in monkeys. Aggression and Violent Behavior. 2000 May 1;5(3):245–54.

42. Maestripieri D. Early experience affects the intergenerational transmission of infant abuse in rhesus monkeys. Proceedings of the National Academy of Sciences. 2005 Jul 5;102(27):9726–9.

43. Maestripieri D, Lindell SG, Higley JD. Intergenerational transmission of maternal behavior in rhesus macaques and its underlying mechanisms. Developmental psychobiology. 2007 Mar;49(2):165–71.

44. Maestripieri D. Neuroendocrine mechanisms underlying the intergenerational transmission of maternal behavior and infant abuse in rhesus macaques. In Hormones and Social Behaviour 2008 May 31 (pp. 121–130). Berlin, Heidelberg: Springer Berlin Heidelberg.

45. Melo L, Pontes AM, da Cruz MM. Infanticide and cannibalism in wild common marmosets. Folia Primatologica. 2003;74(1):48.

46. Miska EA, Ferguson-Smith AC. Transgenerational inheritance: Models and mechanisms of non–DNA sequence–based inheritance. Science. 2016 Oct 7;354(6308):59–63.

47. Osman M, Olkun A, Maldonado AM, Lopez‐Tremoleda J, Sanchez‐Perea N, Paredes UM. Parentally deprived juvenile Owl monkeys suffer from long‐term high infection rates but not from altered hair cortisol concentrations nor from stereotypic behaviours. Journal of Medical Primatology. 2021 Dec;50(6):306–12.

48. Ozaki T, Ohshima T, Miura H, Sato K. Enhancement of miRNA-223-5p and miRNA-30b-5p expression in the saliva of abused children. Dental Journal of Iwate Medical University. 2022 Sep 14;47(1):51–61.

49. Perez G, Barber GP, Benet-Pages A, Casper J, Clawson H, Diekhans M, Fischer C, Gonzalez JN, Hinrichs AS, Lee CM, Nassar LR. The UCSC Genome Browser database: 2025 update. Nucleic Acids Research. 2025 Jan 6;53(D1):D1243–9.

50. Ponnusamy V, Ip RTH, Mohamed MAEK, Clarke P, Wozniak E, Mein C, Schwendimann L, Barlas A, Chisholm P, Chakkarapani E, Michael-Titus AT, Gressens P, Yip PK, Shah DK. Neuronal let-7b-5p acts through the Hippo-YAP pathway in neonatal encephalopathy. Commun Biol. 2021 Sep 30;4(1):1143.

51. Porton I, Niebruegge K. The changing role of hand rearing in zoo-based primate breeding programs. InNursery rearing of nonhuman primates in the 21st century 2006 (pp. 21–31). Boston, MA: Springer US.

52. Prescott MJ, Nixon ME, Farningham DA, Naiken S, Griffiths MA. Laboratory macaques: When to wean?. Applied Animal Behaviour Science. 2012 Mar 1;137(3-4):194–207.

53. Pryce CR, Dettling AC, Spengler M, Schnell CR, Feldon J. Deprivation of parenting disrupts development of homeostatic and reward systems in marmoset monkey offspring. Biological psychiatry. 2004 Jul 15;56(2):72–9.

54. Rechavi O, Lev I. Principles of Transgenerational Small RNA Inheritance in Caenorhabditis elegans. Curr Biol. 2017 Jul 24;27(14):R720–R730.

55. Robertson S, Diver LA, Alvarez-Madrazo S, Livie C, Ejaz A, Fraser R, Connell JM, MacKenzie SM, Davies E. Regulation of corticosteroidogenic genes by microRNAs. International Journal of Endocrinology. 2017;2017(1):2021903.

56. Rogers J. The behavioral genetics of nonhuman primates: status and prospects. American journal of physical anthropology. 2018 Feb;165:23–36.

57. Ryan S, Thompson SD, Roth AM, Gold KC. Effects of hand‐rearing on the reproductive success of western lowland gorillas in North America. Zoo Biology. 2002;21(4):389–401.

58. Sackett GP, Ruppenthal G, Elias K, editors. Nursery rearing of nonhuman primates in the 21st century. Springer Science & Business Media; 2010 May 10.

59. Sanchez MM. The impact of early adverse care on HPA axis development: nonhuman primate models. Hormones and behavior. 2006 Nov 1;50(4):623–31.

60. Sanchez MM, McCormack KM, Maestripieri D. Ethological case study: Infant abuse in rhesus macaques. Formative experiences: The interaction of caregiving, culture, and developmental psychobiology. 2010 Apr 7:224–37.

61. Sánchez N, Galvéz H, Montoya E, Gozalo A. Mortalidad en crías de Aotus sp.(Primates: Cebidae) en cautiverio: una limitante para estudios biomédicos con modelos animales. Revista peruana de medicina experimental y salud pública. 2006 Jul;23(3):221–4.

62. Schino G, Troisi A. Neonatal abandonment in Japanese macaques. American Journal of Physical Anthropology: The Official Publication of the American Association of Physical Anthropologists. 2005 Apr;126(4):447–52.

63. Smith-Vikos T, Slack FJ. MicroRNAs and their roles in aging. Journal of cell science. 2012 Jan 1;125(1):7–17.

64. Sproul Bassett AM, Wood EK, Lindell SG, Schwandt ML, Barr CS, Suomi SJ, Higley JD. Intergenerational effects of mother’s early rearing experience on offspring treatment and socioemotional development. Developmental psychobiology. 2020 Nov;62(7):920–31.

65. Suomi SJ. BIOLOGICAL, MATERNAL, AND LIFE STYLE. East-West Life Expectancy Gap in Europe: Environmental and Non-Environmental Determinants. 1996 Aug 31;19:133.

66. Suomi SJ. Aggression and social behaviour in rhesus monkeys. In Molecular Mechanisms Influencing Aggressive Behaviours: Novartis Foundation Symposium 268 2005 Jul 28 (pp. 216–226). Chichester, UK: John Wiley & Sons, Ltd.

67. Trivers RL. Parent-offspring conflict. American zoologist. 1974 Feb 1;14(1):249–64.

68. Veshkini A, Hammon HM, Lazzari B, Vogel L, Gnott M, Tröscher A, Vendramin V, Sadri H, Sauerwein H, Ceciliani F. Investigating circulating miRNA in transition dairy cows: What miRNAomics tells about metabolic adaptation. Frontiers in Genetics. 2022 Aug 23;13:946211.

69. Yehuda R, Daskalakis NP, Bierer LM, Bader HN, Klengel T, Holsboer F, Binder EB. Holocaust exposure induced intergenerational effects on FKBP5 methylation. Biological psychiatry. 2016 Sep 1;80(5):372–80.

70. Yu F, Wang X, Zhao H, Hao Y, Wang W. Decreased Serum miR-1296 may Serve as an Early Biomarker for the Diagnosis of Non-Alcoholic Fatty Liver Disease. Clinical laboratory. 2019 Oct 1;65(10).

71. Zipple MN, Altmann J, Campos FA, Cords M, Fedigan LM, Lawler RR, Lonsdorf EV, Perry S, Pusey AE, Stoinski TS, Strier KB. Maternal death and offspring fitness in multiple wild primates. Proceedings of the National Academy of Sciences. 2021 Jan 5;118(1):e2015317118.

