## Supplementary material for "Maternal rejecting of newborns is epigenetic, intergenerationally transmitted and associated with altered miRNAs expression in owl monkeys": (Supplementary material 1)

**Supplementary material 1.** Rejection often occurs first births, but some females carry on rejecting afterwards throughout their lives. In graph, orange bars represent counts of rejection by birth order.

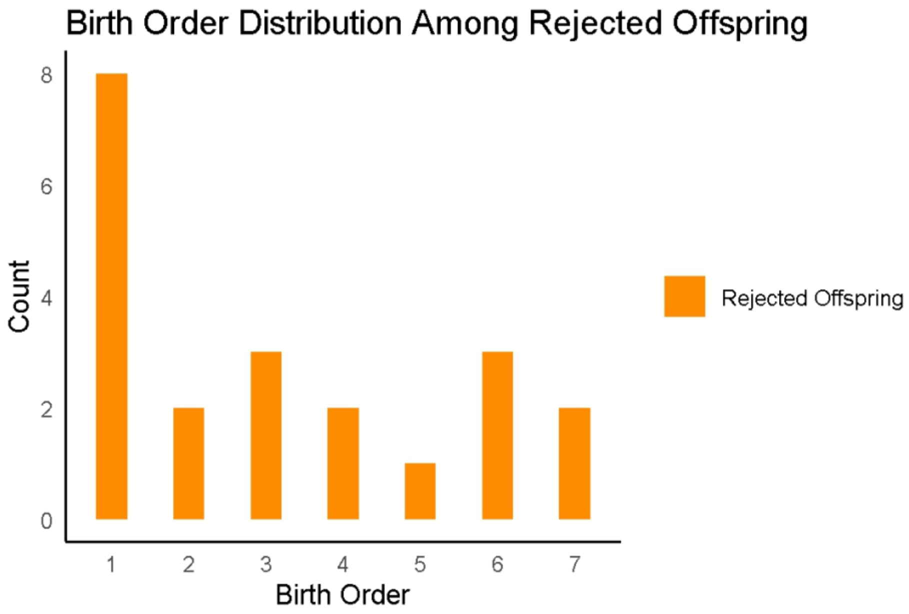
