## Supplementary material for "Maternal rejecting of newborns is epigenetic, intergenerationally transmitted and associated with altered miRNAs expression in owl monkeys": (Supplementary material 2)

**Supplementary material 2.** Some rejected-rejector offspring live lives significantly longer than population average of 8.5-9 years (Farinha & Clausi et al., 2024). In the graph, dots represent ae at death of F1 of rejected-rejectors. In grey those near average lifespan, red circles highlight extended lifespans.

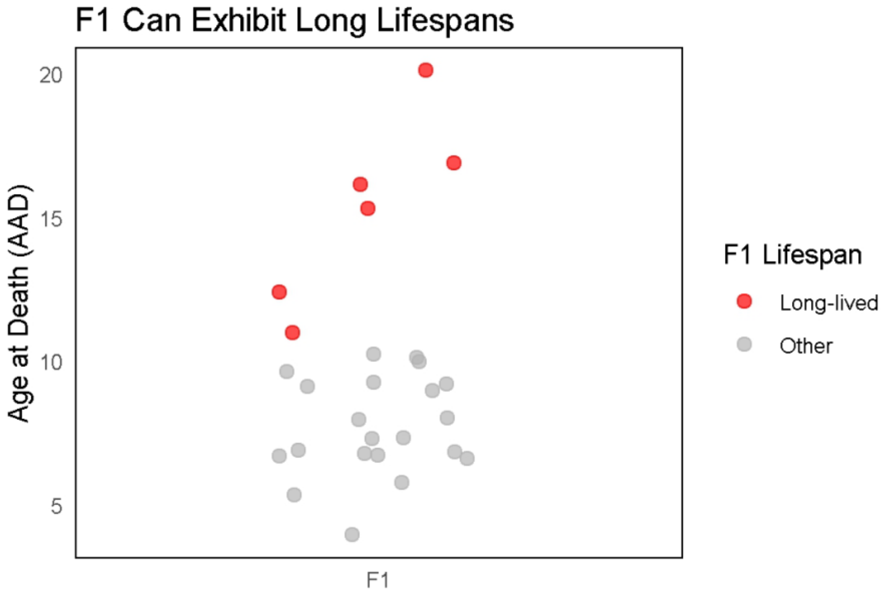
